## Supplementary figures legends for "A single master regulator controls asexual cell cycle division patterns in *Toxoplasma gondii*"

*Supplementary figure legends*

**Figure S1**. Confocal imaging demonstrating the expression of TgAP2IX-5 protein during different stages of the tachyzoite asexual cell cycle using anti-TgCentrin1 as a marker of the cell cycle. (**A**) TgAP2IX-5 expression during the G1/S phase. Expression of TgAP2IX-5 is indicated in red (HA-tag) and TgCentrin1 is indicated in green. DAPI was used to stain the nucleus. Scale bar is indicated at the lower right side of each image. (**B**) TgAP2IX-5 expression during the S/M phase. TgCentrin1 is indicated in green. DAPI was used to stain the nucleus. Scale bar is indicated at the lower right side of each image. (**C**) TgAP2IX-5 expression of parasites within two separate vacuoles at different stages of the cell cycle G1/S and G1. TgAP2IX-5 is indicated in red. TgCentrin1 is indicated in green. DAPI was used to stain the nucleus. Scale bar is indicated at the lower right side of each image.

**Figure S2.** (**A**) Diagram showing the strategy used to generate *Toxoplasma gondii* parasites expressing TgAP2IX-5 tagged with AID-HA-HXGPRT at the C terminus. (**B**) PCR confirming the insertion of AID-HA-HXGPRT insert at the endogenous locus coding for TgAP2IX-5.

**Figure S3.** (**A**) Plaque assay for the parental and iKD TgAP2IX-5 strains in the presence of auxin treatment for 7 days. (**B**) Bar graph representing nucleus per parasite counts for parental and iKD TgAP2IX-5 strains in the absence and presence of 3 hours of auxin treatment. A Student's t-test was performed to compare between the mean percentage of multinucleated parasites between the control (Parental -auxin) and the iKD TgAP2IX-5 mutant strain in the presence of auxin, ** $P < 0.01$; mean ± s.d. (n=3). (**C**) Bar graph representing nucleus per parasite counts for the parental and iKD TgAP2IX-5 strains in the absence and presence of 6 hours of auxin treatment. A Student's t-test was performed to compare between the mean percentage of multinucleated parasites between the control (Parental -auxin) and the iKD TgAP2IX-5 mutant strain in the presence of auxin, ** $P < 0.01$; mean ± s.d. (n=3).

**Figure S4**. (**A**) Confocal imaging of iKD TgAP2IX-5 labelled with TgEno2 (red) and TgISP1 (green) in the presence and absence of auxin treatment. DAPI was used to stain the nucleus. Scale bar is indicated at the lower right side of each image. (**B**) Bar graph representing the percentage of daughter parasite formation in the absence and presence of 6 hours of auxin treatment using TgISP1labelling, A Student's t-test was performed, **P<0.01; mean ± s.d. (n=3).

**Figure S5.** (**A**) Confocal imaging of iKD TgAP2IX-5 labelled with TgChromo1 and TgCentrin1. TgChromo1 is indicated in red. TgCentrin1 is indicated in green. DAPI was used to stain the nucleus. Scale bar is indicated at the lower right side of each image. (**B**) Bar graph representing TgCentrin1: nucleus ratio using the parental and iKD TgAP2IX-5 strains in the absence and presence of auxin treatment for 6 hours. A Student's t-test was performed, **P <0.01; *P < 0.05; mean ±s.d. (n=3). (**C**) Bar graph representing TgChromo1: nucleus ratio using the parental and iKD TgAP2IX-5 strains in the absence and presence of auxin treatment for 6 hours (n=3). A Student's t-test was performed, ***P < 0.001; **P < 0.01; mean ±s.d. (n=3).

**Figure S6.** Effect of TgAP2IX-5 on TgSFA2 and mitochondrion replication in the presence of overnight auxin treatment. **(A)** Confocal imaging of iKD TgAP2IX-5 TgSFA2-myc. TgSFA2 is labelled in red and the nucleus is stained with DAPI. Scale bar is indicated at the lower right side of each image. **(B)** Bar graph representing the ratio of TgSFA2: nucleus in the absence and presence of overnight auxin treatment. A Student's t-test was performed, **P < 0.01; mean ± s.d. (n=3). **(C)** Bar graph representing the ratio of mitochondria: nucleus in the absence and presence of overnight auxin treatment. ***P < 0.001; mean ± s.d. (n=3).

**Figure S7. (A)** Heat map of the cell cycle expression profile for all individual transcripts that are upregulated in the iKD TgAP2IX-5 strain in the presence of 6 hours of auxin treatment. Scale of expression is color-coded with highly expressed genes in orange and less expressed genes in blue according to cell cycle phase indicated at the bottom of the heat map (S-M-C-G1). The cell cycle phases are represented at the bottom as well as the timing when budding occurs. **(B)** Pie chart representing the distribution of the putative localization, according to Barylyuk *et al.*, of the proteins encoded by downregulated transcripts. **(C)** Pie chart representing the localization of downregulated transcripts compared to the whole proteome localization.

**Figure S8.** ChIP-seq data analysis represented by peaks targeting the promoter of several genes. (**A**) MACS2 generated track and individual ChIP tracks (background subtracted) representing the direct targeting of TgAP2IX-5 to the promoter of downregulated gene TgIMC1 and TgIMC4. (**B**) MACS2 track representing the direct targeting of TgAP2IX-5 to the promoter of downregulated gene TgIMC3. **(C)** MACS2 track representing the direct targeting of TgAP2IX-5 to the promoter of downregulated gene TgAP2XII-9. (**D**) ChIP-seq data peak indicated by blue box demonstrating the targeting of TgAP2IX-5 towards upregulated gene TgAP2IX-5 (boxed). (**E**) Heat map representing the cell cycle expression of all individual transcripts that are targeted by TgAP2IX-5 based on ChIP-seq analysis. Phases of the cell cycle are indicated at the bottom of the figure. The approximate timing for the budding cycle is indicated at the bottom.

**Figure S9.** (**A**) Bar graph representing the percentage of daughter parasite formation in the absence and presence of overnight auxin treatment using TgIMC29 labelling, A Student's t-test was performed, $*P < 0.05$; mean ± s.d. (n=3). (**B**) Heat map of 8 individual TgApiAP2 TF transcripts that are downregulated during 6 hours of TgAP2IX-5 depletion. Cell cycle phases are indicated at the lower bottom (S-M-C-G1). Downregulated ApiAP2 TF that are directly bound to promoters are underlined in green.

**Figure S10.** (**A**) TgAP2IX-5 iKD complementation schematic representation demonstrating the strategy used for generating the complemented TgAP2IX-5 iKD strain by targeting the UPRT locus and replacing it with exogenous myc-tagged TgAP2IX-5 under the control of its own specific promoter. (**B**) Bar graph representing the expression of iKD TgAP2IX-5 using anti-HA antibody and anti-myc antibody as a negative control.

**Figure S11.** (**A**) Immunofluorescence assays of iKD TgAP2IX-5 parasites before auxin washout and after auxin washout for a duration of 4 hrs. TgIMC3 is labelled in red. TgISP1 is labelled in green. DAPI was used to stain the nucleus. Scale bar is indicated at the lower right side of each image.

**Video S1.** Time-lapse video of iKD TgAP2IX-5-TgIMC3-mCherry mutant after being treated with auxin for 16 hours subsequently followed by auxin washout for 3.9 hours consisting of a compilation of images taken every 13 minutes using a spinning-disc microscope.

**Supplementary Table legends**

**Table S1:** Oligonucleotides used in this study.

**Table S2:** RNASeq results. Summary of all genes up or downregulated with a fold change above (-1 or 1 of $Log_2$ fold change) and a FDR of 0.05.

**Table S3: ChIP-seq results.** Gene IDs of the promoters bound by TgAP2IX-5 as identified by MACS2 with a FDR of 0.05.
