## Supplementary figures for "A single master regulator controls asexual cell cycle division patterns in *Toxoplasma gondii*"

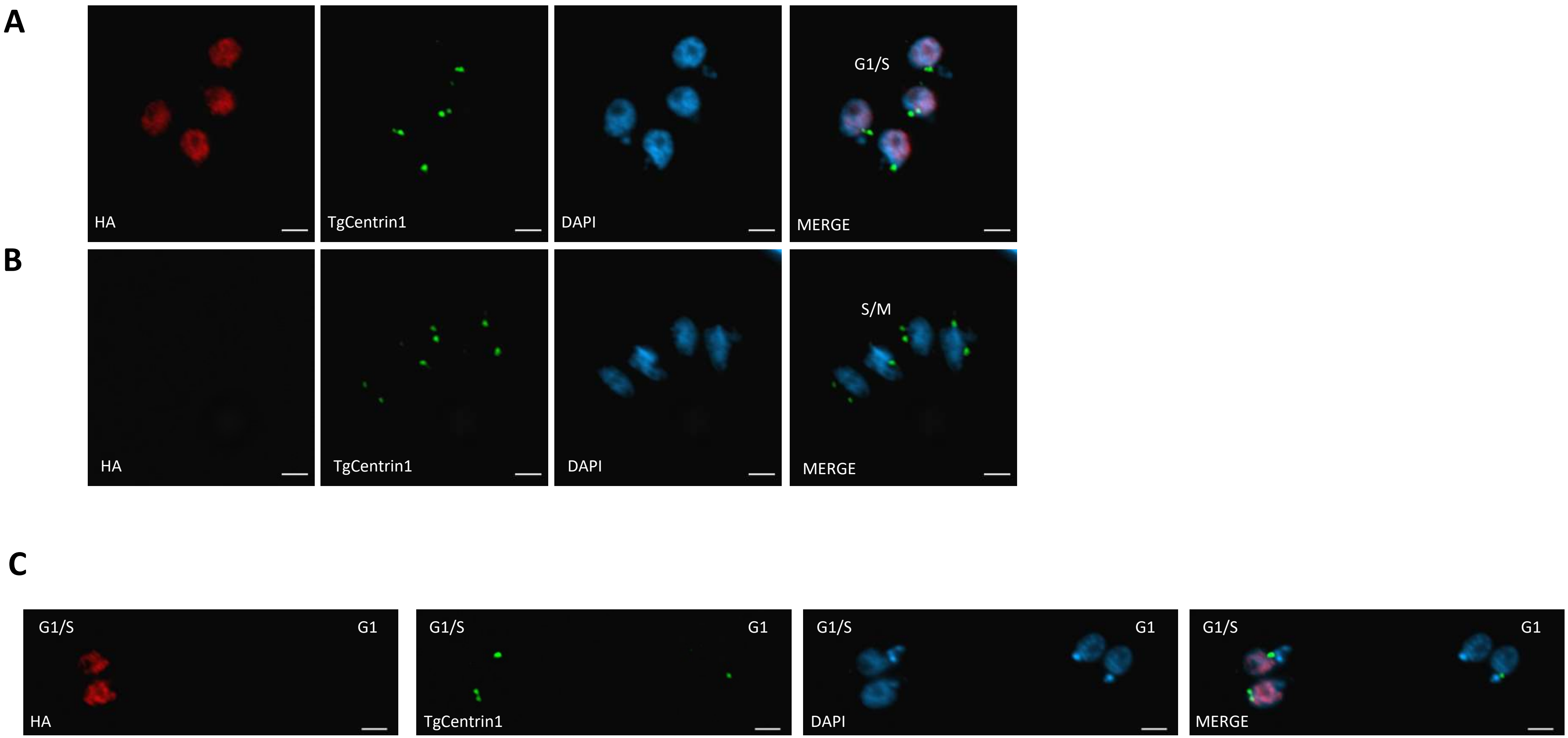
A
HA
TgCentrin1
DAPI
G1/S
MERGE
B
HA
TgCentrin1
DAPI
S/M
MERGE
C
G1/S
G1
HA
G1/S
G1
TgCentrin1
G1/S
G1
DAPI
G1/S
G1
MERGE

Figure S1

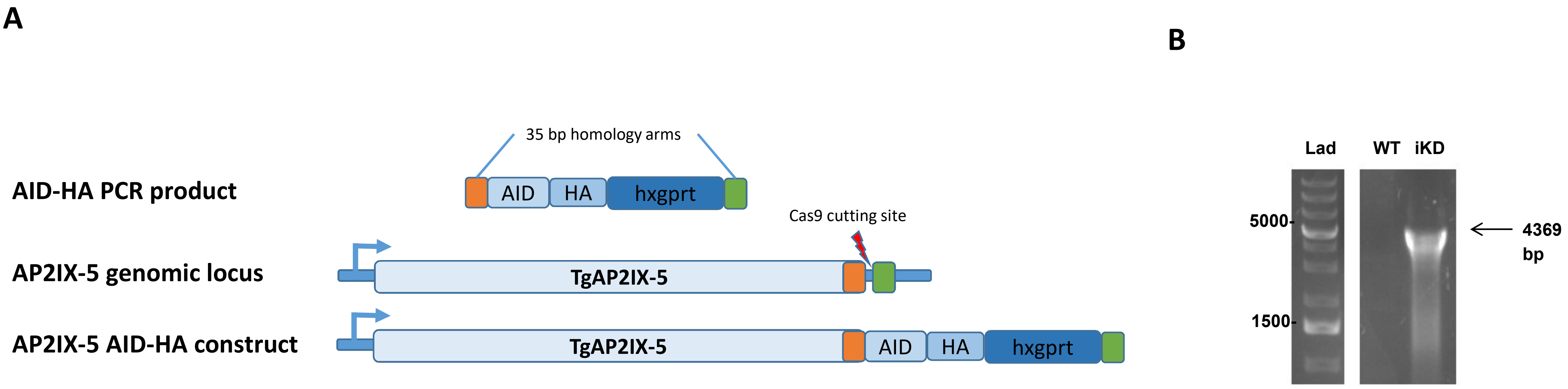
A
35 bp homology arms
AID-HA PCR product
AID
HA
hxgprt
Cas9 cutting site
AP2IX-5 genomic locus
TgAP2IX-5
AP2IX-5 AID-HA construct
TgAP2IX-5
AID
HA
hxgprt
B
Lad
WT
iKD
5000-
1500-
4369
bp

Figure S2

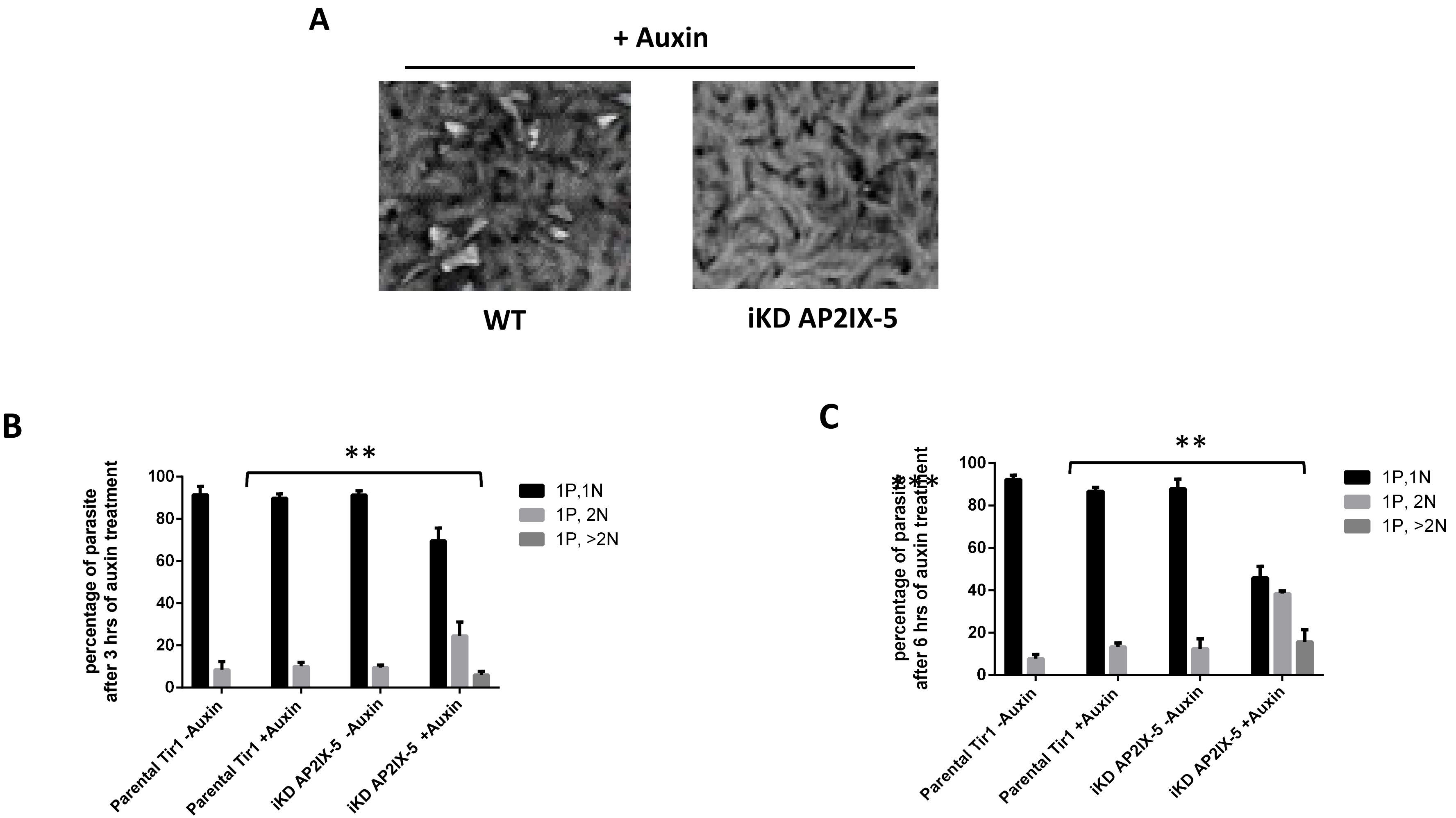
A
+ Auxin
WT
iKD AP2IX-5
B
**
percentage of parasite after 3 hrs of auxin treatment
100
80
60
40
20
0
1P,1N
1P, 2N
1P, >2N
Parental Tir1 -Auxin
Parental Tir1 +Auxin
iKD AP2IX-5 -Auxin
iKD AP2IX-5 +Auxin
C
**
percentage of parasite after 6 hrs of auxin treatment
100
80
60
40
20
0
1P,1N
1P, 2N
1P, >2N
Parental Tir1 -Auxin
Parental Tir1 +Auxin
iKD AP2IX-5 -Auxin
iKD AP2IX-5 +Auxin

Figure S3

A

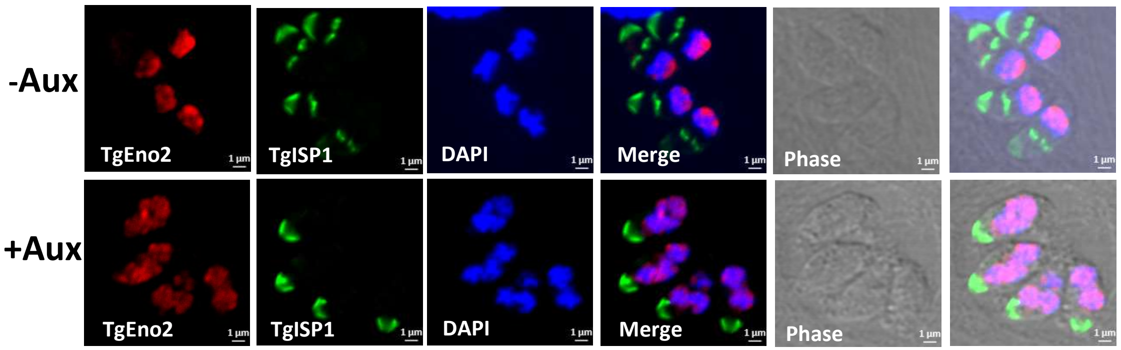

-Aux
+Aux
TgEno2
TgISP1
DAPI
Merge
Phase
1 µm


B

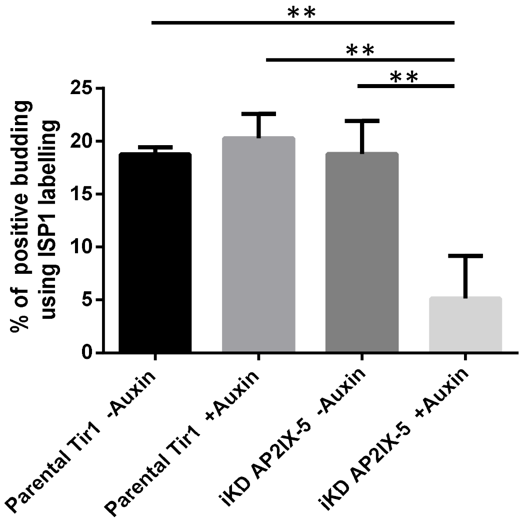

% of positive budding using ISP1 labelling
25
20
15
10
5
0
**
**
**
Parental Tir1 -Auxin
Parental Tir1 +Auxin
iKD AP2IX-5 -Auxin
iKD AP2IX-5 +Auxin


Figure S4

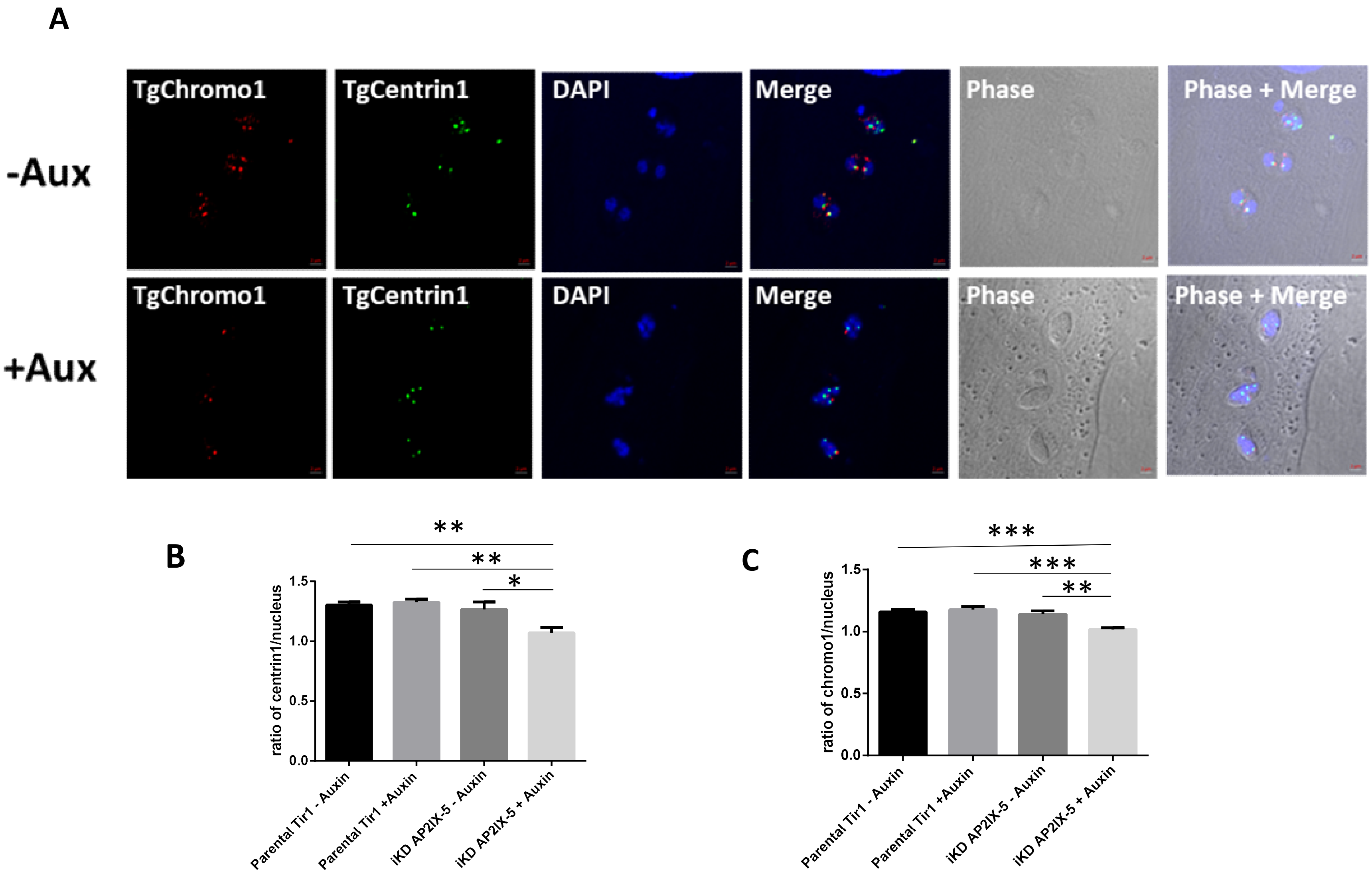
A
-Aux
+Aux
TgChromo1
TgCentrin1
DAPI
Merge
Phase
Phase + Merge
B
ratio of centrin1/nucleus
0.0
0.5
1.0
1.5
**
**
*
Parental Tir1 - Auxin
Parental Tir1 +Auxin
iKD AP2IX-5 - Auxin
iKD AP2IX-5 + Auxin
C
ratio of chromo1/nucleus
***
***
**

Figure S5

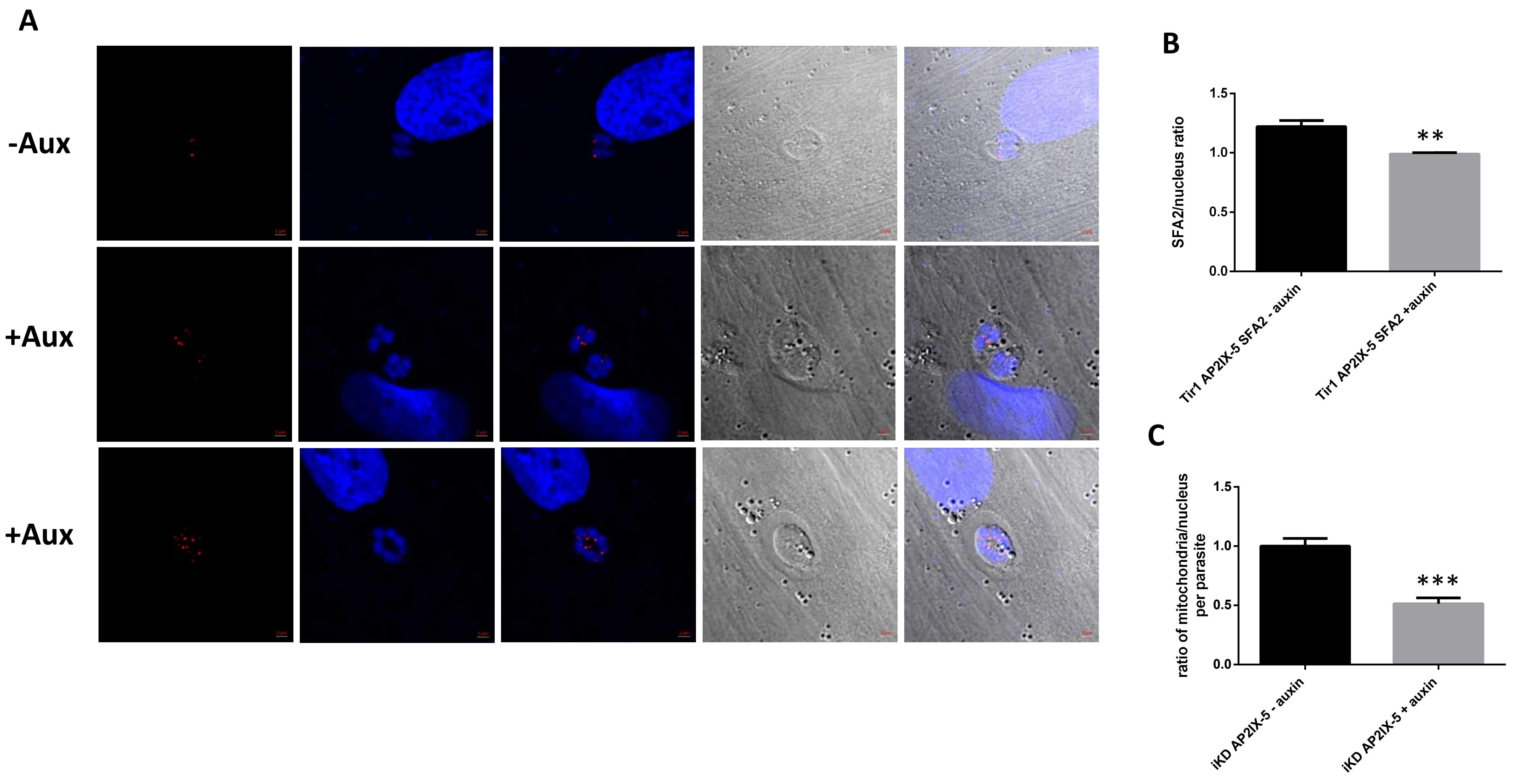
A
-Aux
+Aux
+Aux
B
SFA2/nucleus ratio
1.5
1.0
0.5
0.0
**
Tir1 AP2IX-5 SFA2 - auxin
Tir1 AP2IX-5 SFA2 +auxin
C
ratio of mitochondria/nucleus per parasite
1.5
1.0
0.5
0.0
***
iKD AP2IX-5 - auxin
iKD AP2IX-5 + auxin

Figure S6

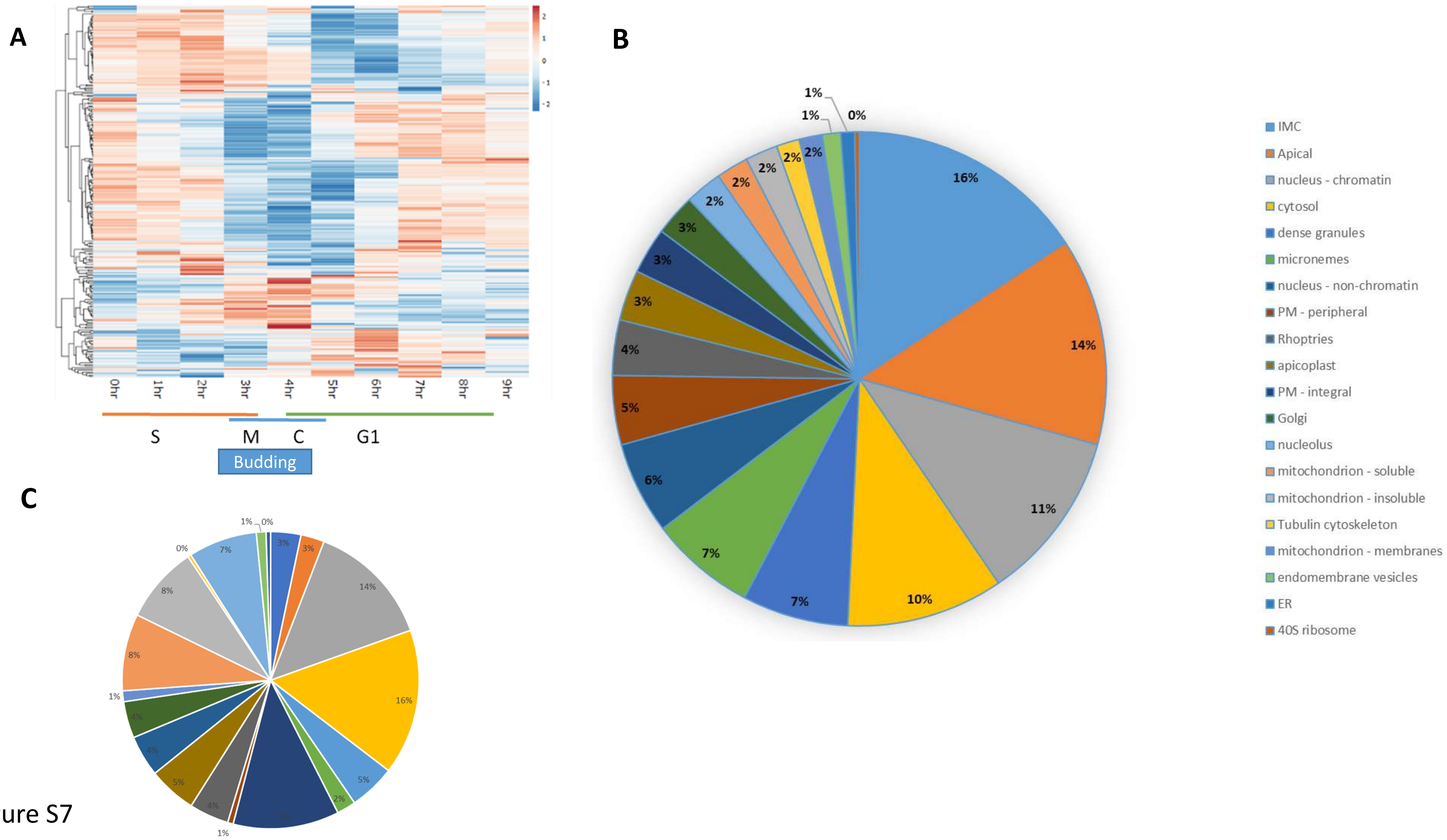
A
0hr
1hr
2hr
3hr
4hr
5hr
6hr
7hr
8hr
9hr
S
M
C
G1
Budding
B
16%
14%
11%
10%
7%
7%
6%
5%
4%
3%
3%
3%
2%
2%
2%
2%
2%
1%
1%
0%
IMC
Apical
nucleus - chromatin
cytosol
dense granules
micronemes
nucleus - non-chromatin
PM - peripheral
Rhoptries
apicoplast
PM - integral
Golgi
nucleolus
mitochondrion - soluble
mitochondrion - insoluble
Tubulin cytoskeleton
mitochondrion - membranes
endomembrane vesicles
ER
40S ribosome
C

Figure S7

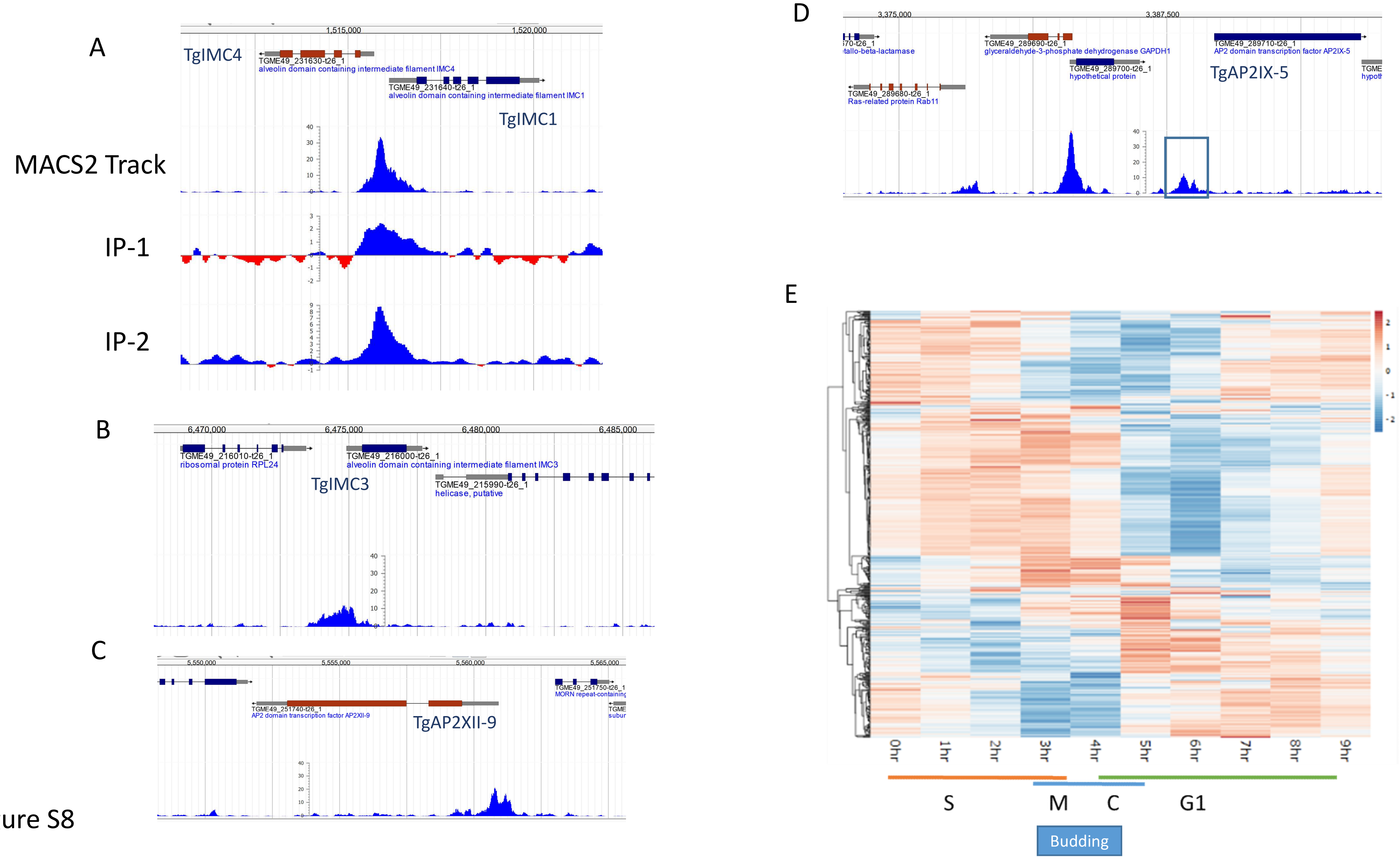
A
TgIMC4
TGME49_231630-t26_1
alveolin domain containing intermediate filament IMC4
TGME49_231640-t26_1
alveolin domain containing intermediate filament IMC1
TgIMC1
MACS2 Track
IP-1
IP-2
B
TGME49_216010-t26_1
ribosomal protein RPL24
TGME49_216000-t26_1
alveolin domain containing intermediate filament IMC3
TgIMC3
TGME49_215990-t26_1
helicase, putative
C
TGME49_251740-t26_1
AP2 domain transcription factor AP2XII-9
TgAP2XII-9
TGME49_251750-t26_1
D
TGME49_289690-t26_1
glyceraldehyde-3-phosphate dehydrogenase GAPDH1
TGME49_289710-t26_1
AP2 domain transcription factor AP2IX-5
TGME49_289700-t26_1
hypothetical protein
TgAP2IX-5
TGME49_289680-t26_1
Ras-related protein Rab11
E
0hr
1hr
2hr
3hr
4hr
5hr
6hr
7hr
8hr
9hr
S
M
C
G1
Budding

Figure S8

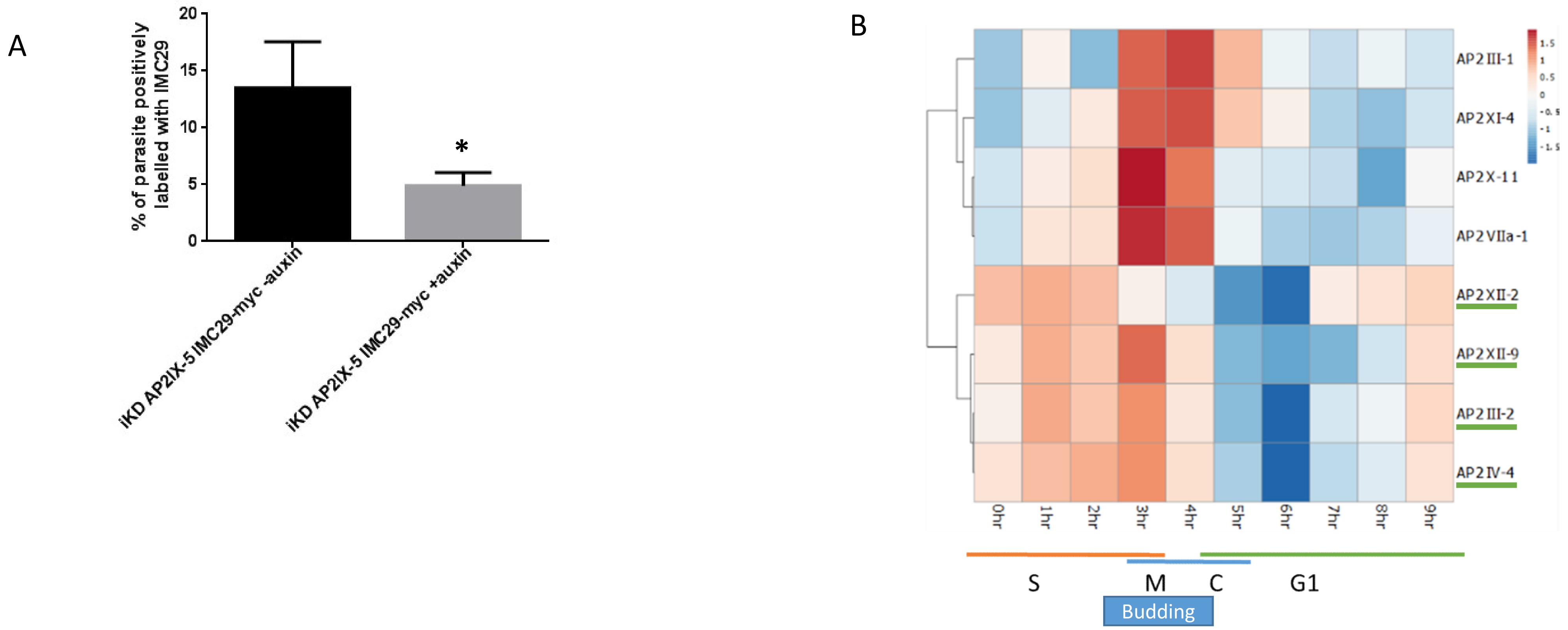
A
% of parasite positively labelled with IMC29
20
15
10
5
0
*
iKD AP2IX-5 IMC29-myc -auxin
iKD AP2IX-5 IMC29-myc +auxin
B
AP2III-1
AP2XI-4
AP2X-11
AP2VIIa-1
AP2XII-2
AP2XII-9
AP2III-2
AP2IV-4
0hr
1hr
2hr
3hr
4hr
5hr
6hr
7hr
8hr
9hr
S
M
C
G1
Budding

Figure S9

A

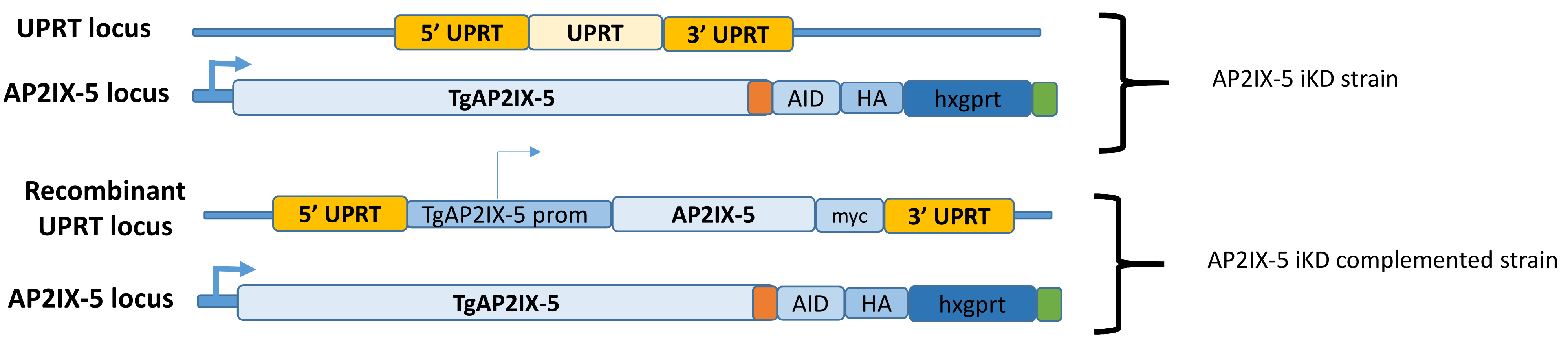


B

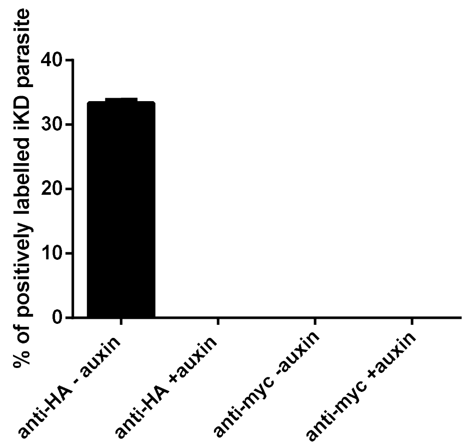


Figure S10

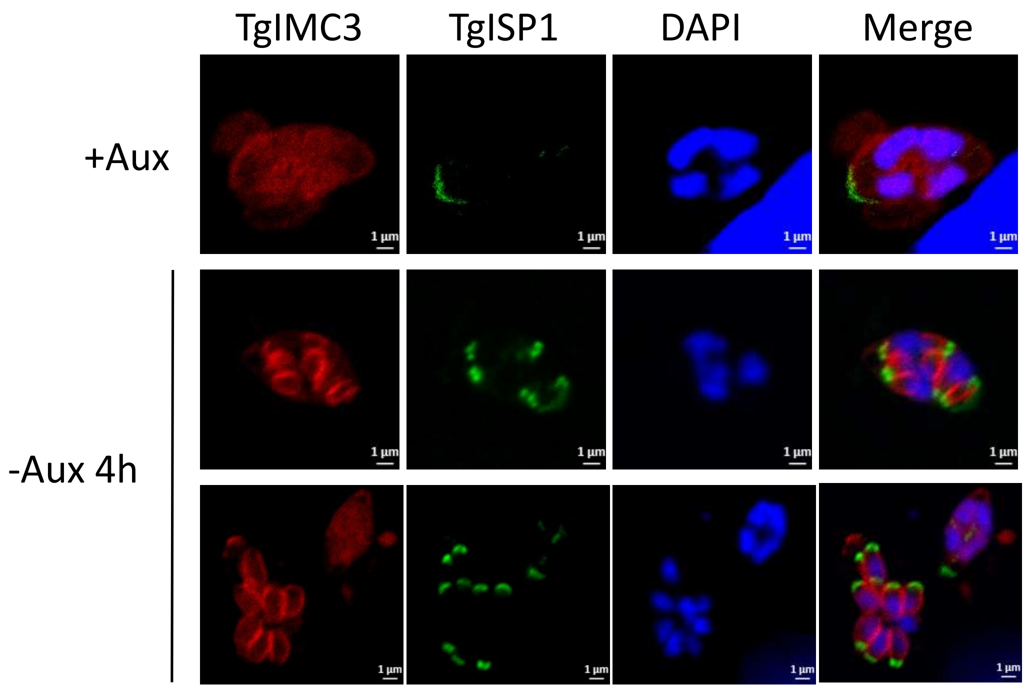


Figure S11

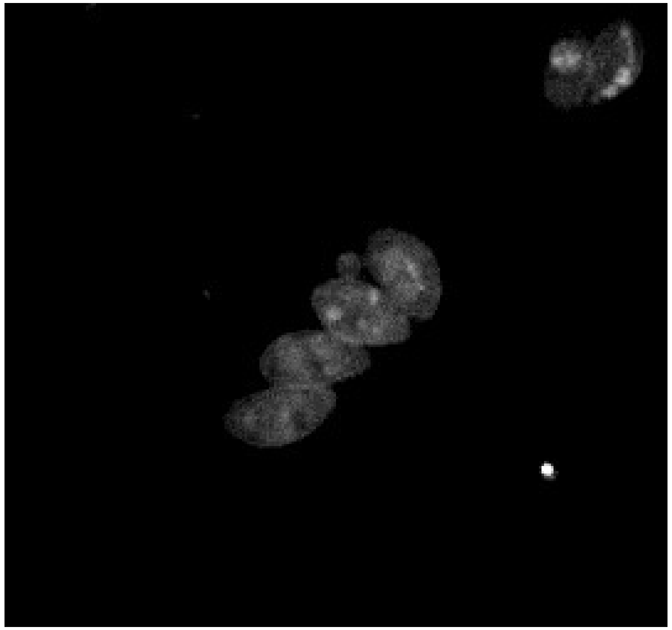

Video S1
